# Transcriptional activity and epigenetic regulation of transposable elements in the symbiotic fungus *Rhizophagus irregularis*

**DOI:** 10.1101/2021.03.30.436303

**Authors:** A Dallaire, BF Manley, M Wilkens, I Bista, C Quan, E Evangelisti, CR Bradshaw, NB Ramakrishna, S Schornack, F Butter, U Paszkowski, EA Miska

## Abstract

Arbuscular mycorrhizal (AM) fungi form mutualistic relationships with most land plant species and have long been considered as ancient asexuals. Long-term clonal evolution would be remarkable for a eukaryotic lineage and suggests the importance of alternative mechanisms to promote genetic variability facilitating adaptation. Here, we assessed the potential of transposable elements (TEs) for generating genomic diversity. The dynamic expression of TEs during *Rhizophagus irregularis* spore development suggests ongoing TE activity. We find *Mutator-like* elements located near genes belonging to highly expanded gene families. Characterising the epigenomic status of *R. irregularis* provides evidence of DNA methylation and small RNA production occurring at TE loci. Our results support a potential role for TEs in shaping the genome, and roles for DNA methylation and small RNA-mediated silencing in regulating TEs. A well-controlled balance between TE activity and repression may therefore contribute to genome evolution in AM fungi.

## Introduction

The AM symbiosis is hundreds of millions of years old, and a majority of the world’s plant species are hosts to AM fungi (1). As such, these fungi exist in a wide range of environments and can even engage in symbioses with multiple plant species simultaneously. The complex life cycles of AM fungi suggest a requirement for strong developmental and phenotypic plasticity. However, genetically distinct strains created by meiosis have never been reported (2, 3) and, although they carry meiosis-related genes (4), evidence of sexual reproduction is lacking (5). This has led to the hypothesis that AM fungi are ancient asexual organisms, which raises a key question on how these fungi were able to diversify their gene inventory and fill such varied ecological niches.

Genome assemblies are available for a number of AM fungal species, including *R. irregularis* (6, 7, 8), *R. diaphanus, R. cerebriforme, Gigaspora rosea* (9)*, Diversispora epigaea* (10), and *G. maragarita* 11). Genomic analyses of these species have revealed contents of repetitive sequences ranging from 23 to 43% (9, 12). These repeats consist of transposable elements (TE) and expanded gene families that occasionally form tandemly repeated arrays of duplicate genes (6, 7, 8, 9, 13, 14). Expanded family genes are either orphans (no significant homologs can be identified), or contain protein domains related to signalling and RNA interference (RNAi), such as kinase, BTB/POZ (Broad-Complex, Tramtrack and Bric a brac/Pox virus and Zinc finger) domains, Sel1 tetratricopeptide repeats, Kelch-repeats and P-element Induced WImpy testis (PIWI) domains. High copy number genes are present in variable numbers in AMF isolates and although their evolutionary origins and functions are unknown, they have been proposed to play roles in perception and interaction with the environment (15).

Transposons are repetitive DNA sequences that colonise genomes and generate intra- and inter-specific genetic variability. They are able to move and replicate within genomes and can increase genomic plasticity by promoting chromosomal rearrangements and compartmentalisation, duplicating or deleting genes, and altering gene expression (16). As their accumulation can also have deleterious effects on their host genome, eukaryotes have developed defence mechanisms to control TE proliferation. Three mechanisms of defence have been described in fungi: repeat-induced point mutation (RIP)(17), DNA cytosine methylation (18), and small RNA (sRNA)-mediated gene silencing (19). Signatures of RIP have not been detected in the AM fungus *G. margarita* or in species of the Mucoromycotina, a sister subphylum to the Glomeromycotina to which AM fungi belong (20). However, the presence of DNA cytosine methyltransferases and sRNA pathway genes encoded in AM genomes suggests their role in TE silencing.

The extent to which TEs have shaped the genome evolution of AM fungi is still largely unknown, as are the mechanisms by which these species keep TEs under control. In this study, we investigate the organisation of TEs, DNA methylation and sRNAs on a global genome level in the model AM fungus *R. irregularis*. We bring together single-molecule DNA methylation detection, sRNA/transcriptome sequencing and proteomics to uncover fundamental characteristics of epigenetic mechanisms in AM fungi. This study reveals signs of TE activity in *R. irregularis* and uncovers a potential role for *Mutator-like* elements in expanding specific gene families. We provide evidence for DNA methylation and small RNA-mediated TE regulation.

## Highlights

- Evidence of recent or ongoing TE activity in *R. irregularis*.
- The location of high copy number genes is linked to TEs.
- A potential role for *Mutator-like* elements in expanding specific gene families, notably RNAi pathway genes.
- First methylome analysis in the Glomeromycotina subphylum.
- DNA methylation and small RNA production at TE loci.

## Results

### The TE landscape of *R. irregularis*

In order to understand how TEs have shaped the genome of *R. irregularis*, we first generated a new TE annotation of the published *R. irregularis* genome (8), which was produced using long-read sequencing. Our annotation revealed a 47% repeat coverage, which is consistent with the previous report by Maeda et al (8) (Figure S1AB). Of these repeats, 12% could be classified into TE families, which is within the range of what is typically observed in fungi (0.2-30% (21)), but lower compared to the obligate biotroph pathogen *Blumeria graminis* (∼45% 22)). The genome of *R. irregularis* contains many DNA transposons and retrotransposons such as Gypsy LTRs, LINE R1, Hobo and Tc1 (Figure S1B). A portion of repetitive sequences occupying 32% of the genome could not be classified into TE families (Figure S1ABC). Repeats overlapping with high copy number (kinase, BTB/POZ, Sel1, Kelch-like) and orphan genes account for some of these unclassified repeats (8% of the genome Figure S1A). The remaining unclassified repeats (occupying 24% of the genome) may be repetitive elements unique to this species. Subsequent analyses were focused on classified TEs. Using divergence analysis based on calculated Kimura distance, we observe two waves of transposon expansion in the *R. irregularis* genome (Figure 1A). Recent expansions include DNA transposons, retrotransposon and helitron activity (Kimura distance 0-1, Figure 1A). More specifically, families which show recent activity include the DNA transposons Maverick, CMC, hAT, MULE and Gypsy retrotransposons (Figure S1C). These TE classes have therefore shaped genome architecture in *R. irregularis*.

**Figure 1.**
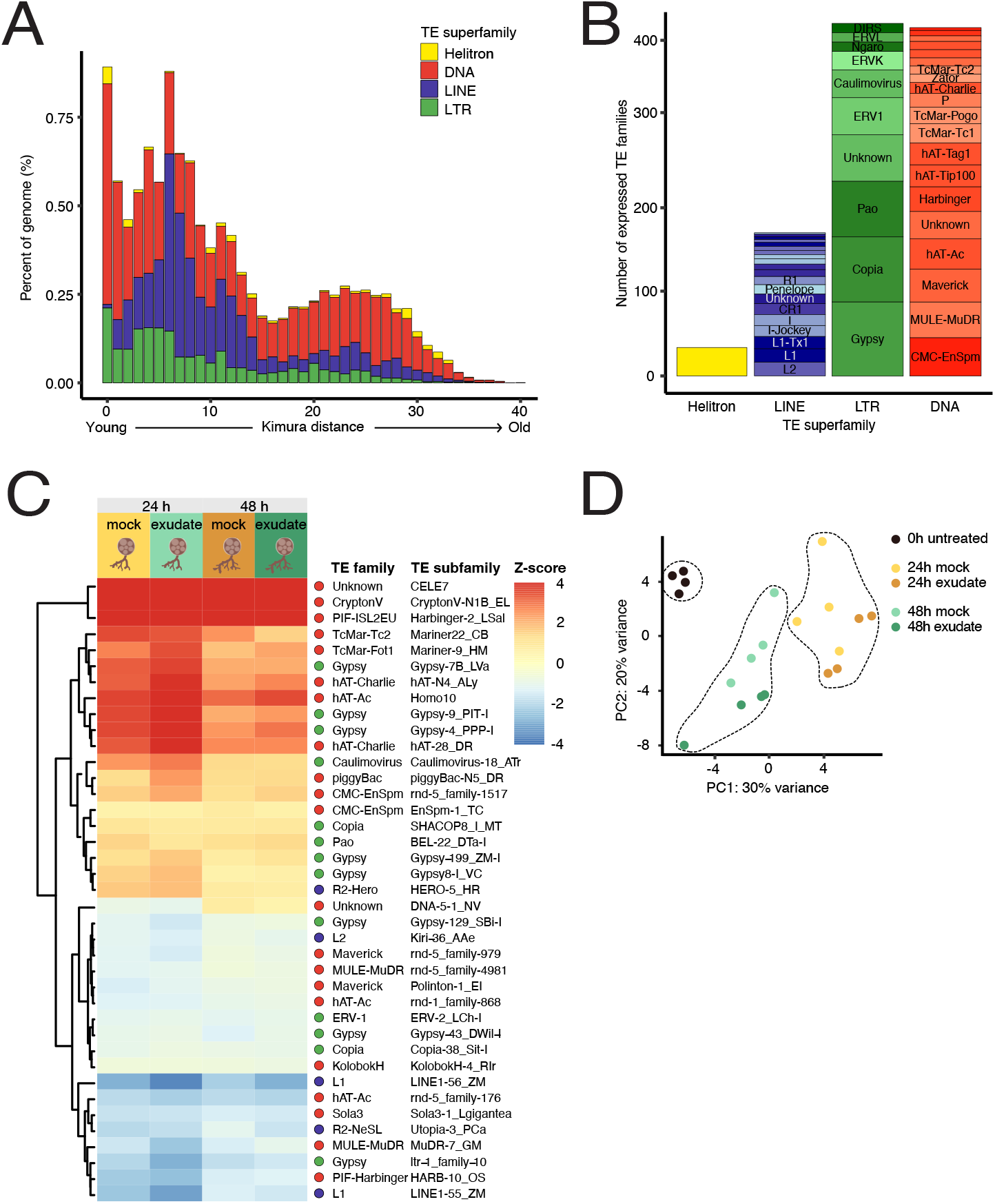
Transposon expression during *R. irregularis* spore development. **A**. Repeat landscapes showing Kimura distance-based copy divergence analysis of TEs in *R. irregularis* genome shown as genome coverage (%) for each TE superfamily plotted against Kimura distance. Clustering was performed according to their Kimura distances of TEs (CpG adjusted *K-value* from 0 to 50). TE copies with a low Kimura distance value have a low divergence from the consensus sequence and may correspond to recent replication events. Sequences with a higher Kimura distance value corresponded to older divergence. Note that we omitted unclassified elements (For repeat landscape including unclassified elements see Figure S1). **B**. Number of expressed transposon subfamilies, grouped superfamilies. **C**. Heatmap and hierarchical clustering of differentially expressed TE subfamilies (|log_2_FC| > 0.5; FDR <0.05) in the spore developmental assay. Five conditions were used in total, a 0h control treatment, 24h mock and rice exudate treatments, and 48h mock and rice exudate treatments (4 replicates per treatment). Expression of 24h and 48h conditions was normalised against expression in the control, 0h condition. **D**. Principal component analysis of TE subfamily expression across all replicates and conditions.

### Expression of TEs in developing *R. irregularis* spores

The process leading to TE activation is not well understood, but transcription is a precondition for their mobility. We measured the expression levels of TE transcripts in a developmental assay where *R. irregularis* spores were exposed to either medium containing rice root-derived exudates or a mock nutrient medium. Spore response to these two conditions was examined at three time points: a 0h untreated time point, 24h and 48h post treatment. An initial global analysis was carried out to establish the full scope of TE and gene expression at any of these time points. Using tools optimised for quantifying highly repetitive sequences, we detected significant expression for members of all TE superfamilies (Figure 1B). The most represented families included Gypsy, Copia, Pao, CMC-EnSpm and MULE-MuDR elements (Figure 1B). We then investigated the expression of TEs at the subfamily level during the spore development assay. While a majority of differentially expressed TE families were DNA transposons, LINEs and LTRs also displayed both up- and down-regulation during the developmental assay (Figure 1C, Table S1). Principal component analysis (PCA) of TE subfamily expression showed that biological replicates formed discrete clusters based on time point but did not cluster well in response to treatment (rice exudates or mock treatment) (Figure 1D). This may indicate that TE subfamily expression dynamics are not dependent on plant-derived compounds under these conditions. We next sought to analyse TE expression with locus-level resolution. We detected 786 individual TEs that overlapped with expressed genes (Table S2). Of those, 232 had significant expression and shorter lengths (Figure S2A), but were excluded from following analyses as their expression could be attributed to expression of the genes they reside in. Individual TEs displayed wide ranges of expression levels, belonged to all TE superfamilies (Figure S2B) and could be categorised into evolutionary divergence bins between 0 and 40 (Figure S2C), suggesting that TEs of all ages are transcribed. Dynamic expression of TEs suggests ongoing activity, however, in order to mobilise, TEs would likely need to be full-length elements. We therefore examined the length and relative age of expressed TEs. TEs with lengths of over 2kb and low Kimura distance were detected (Figure S2D; shaded area). However, due to ambiguous mapping of reads to individual TE copies, we could not validate the expression of full-length copies. Taken together, this data shows fluctuations in TE subfamily expression in germinating spores, which may indicate developmental relaxation of TE silencing during spore development and suggests a potential ongoing TE activity in *R. irregularis*.

### Methylome analysis of spore DNA using single-molecule sequencing

Detection of TE expression led us to hypothesise that epigenetic mechanisms may be regulating TE activity in spores. Cytosine methylation is an important factor in suppression of TE transcription (23). We therefore assessed the role of DNA methylation in regulating TEs by surveying genome-wide 5-methylcytosine (5mC) using Nanopore long-read direct sequencing of DNA extracted from untreated spores. A total of 2,876,042 mCG sites were identified, accounting for 13.8% of total genomic cytosine content. CG site methylation status displayed a strong bimodal distribution, with sites either highly (30.8% of CpGs >0.8) or weakly (60.3% of CpGs <0.2) methylated (Figure 2A). This trend of bimodal CG site methylation is also observed in plants, animals and other fungi (24–27). In fungal species studied so far, it has been found that limited methylation occurs in gene bodies and methylation levels are highest in repeats and transposons (18). To examine whether this was the case in *R. irregularis*, we profiled the levels of mCG in classified TEs (Figure 2A). The median mCG levels were high in most transposon families. However, the MULE-MuDR elements found in our analysis to be down-regulated during spore development (Figure 1C) displayed lower than average median mCG levels (12.5 and 25% respectively, Figure 2B). Low methylation coupled with dynamic expression may indicate that MULEs are still active. Other recently expanded TE classes displayed a high proportion of TE copies with low methylation, including DNA/TcMar, DNA/hAT, LTR/Gypsy and DNA/Maverick elements. The low methylation levels of many TE copies may indicate the absence of transcriptional suppression, or transcriptional suppression through other control layers such as CHG/CHH methylation, histone modification, or small RNA-mediated interference (RNAi). In general, short and evolutionarily older TEs displayed low mCG scores (Figure 2C), suggesting a loss of mCG in old, degenerated TEs, a phenomenon also observed in rice retrotransposons and mammalian LINEs (28, 29). We examined the mCG levels along the length of TEs and in flanking regions and found higher mCG levels within the TE locus than in the immediate upstream and downstream regions (Figure 2D). DNA hypomethylation can lead to TE de-repression if no other restraining mechanism is active. We therefore examined whether low mCG levels could be linked to TE loci expression. Indeed, members of all families of expressed TEs were significantly associated with lower mCG levels compared to their respective non-expressed counterparts (Figure 2E). This data shows that *R. irregularis* TEs are generally highly methylated, but old, short and expressed TEs tend to have lower mCG levels. DNA methylation may therefore be associated with the regulation of TEs during development and over evolutionary time.

**Figure 2.**
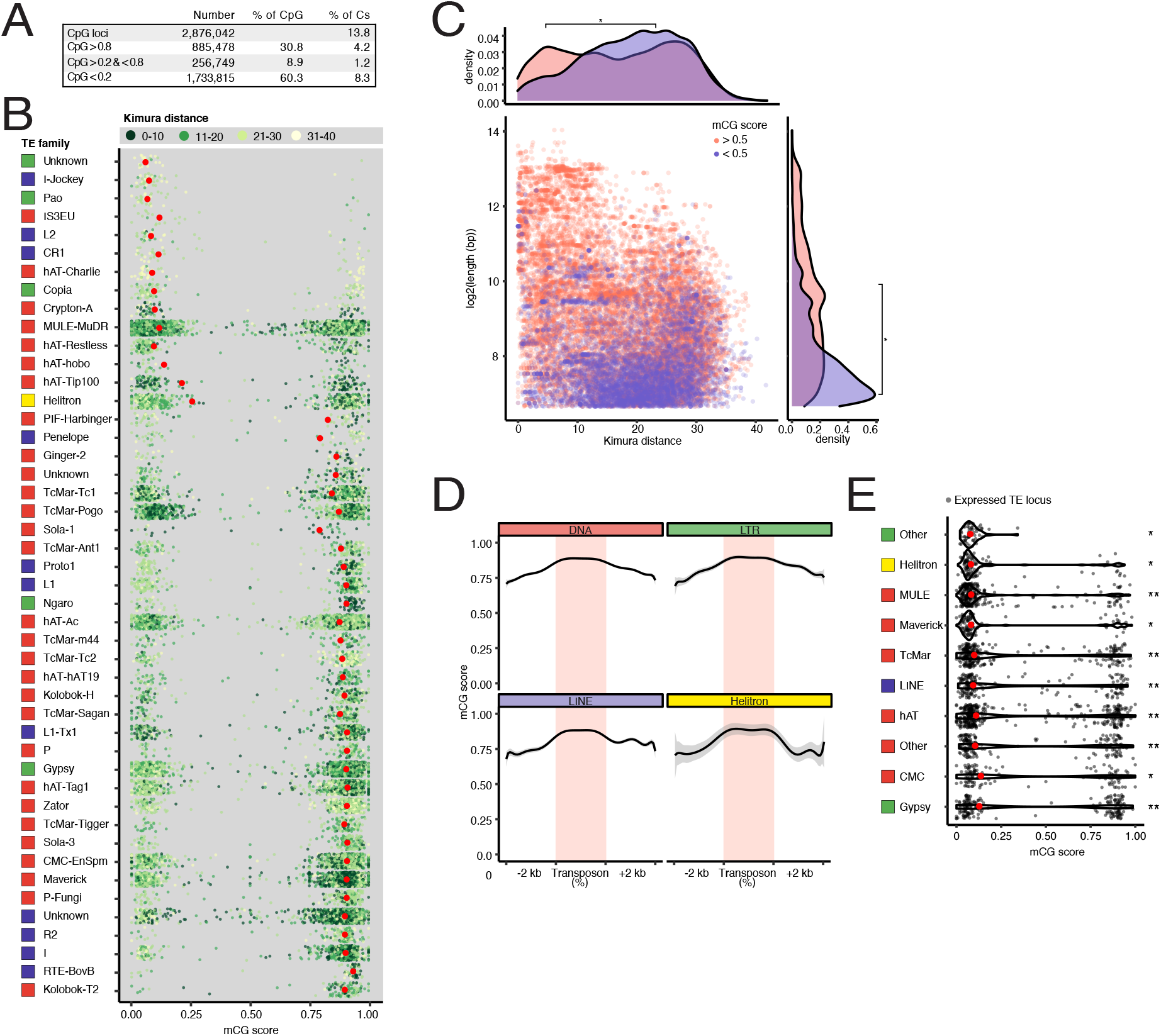
Interplay between transposons and DNA methylation in *R. irregularis* spores. **A**. Absolute and relative proportions of mCG sites in *R. irregularis* spores. **B**. Average methylation (mCG score) of individual TE loci (length >100bp and copy number >20), grouped into TE families. Red dots show the median values and point colour indicate the relative age of each TE expressed as Kimura distance and grouped into bins. **C**. Length of TEs shown in B, relative to divergence expressed as Kimura distance. Point colour indicates high and low mCG score (pink and purple respectively). Density plots depict TE length and divergence for elements belonging to each mCG score category. A Kruskal-Wallis H test was performed to compare the mCG score distributions of lowly and highly methylated TEs. *p-value<2.2E-16. **D**. Metagene plots displaying mCG levels across two groups of TEs, lowly methylated (mCG score < 0.5), and highly methylated (mCG score > 0.5), and their 2-kb upstream and downstream sequences. **E**. mCG scores of expressed (grey) and differentially expressed (green) TEs. Red dots represent the median values of each super-family. Significance was assessed by a Kruskal-Wallis H test comparing the mCG score distribution of expressed TEs to the mCG score distribution of non-expressed TEs of the same class. *Kruskal-wallis p-value <1E-36 >1E-100. **Kruskal-wallis p-value<1E-100.

### High copy number genes are located close to *Mutator-like* elements (MULEs)

The genome of *R. irregularis* is particularly interesting because it contains over 2000 kinases and 30 Argonaute (AGO) genes, the highest numbers of these genes recorded so far in any species (6, 7, 8, 9 13, 14). As all superfamilies of transposons are capable of duplicating genes or gene fragments (30), we hypothesized that TE activity may have caused these gene expansions. We profiled mCG levels in gene bodies and observed that, although the majority of genes displayed very low mCG levels, 18% of all *R. irregularis* genes are highly methylated (Figure 3A). This feature that sets *R. irregularis* apart from other fungi, which typically have low gene body methylation (18). The distribution of mCG across genic regions of highly methylated genes was similar to that seen in TEs (Figure 3B). We wondered what these highly methylated genes could be and, because the genome of *R. irregularis* has a high content of repetitive genes (Figure S1A), we hypothesised that repetitiveness may correlate with transcriptional silencing by DNA methylation. We therefore examined the functional annotation of lowly and highly methylated genes and categorised them based on protein-coding domains: Class A are core, low copy number genes, Class B have no known protein domain (orphan genes), and Class C are high copy number genes with serine/threonine/tyrosine kinase, calmodulin-dependent kinase, BTB/POZ, Sel1-like or Kelch-like domains. Only the most highly amplified gene families with known proteins domains (described in 6, 8) were included in the high copy number class C. The remainder are either genes with transposon-related domains (e.g. reverse transcriptase), or crinkler domains (PFAM PF20147), part of a subfamily of candidate AMF secreted effector proteins (31). In plant pathogenic fungi, such candidate secreted effector genes tend to be located in TErich genomic islands (32), and some have co-evolved with particular TEs (33, 34). We found that 61.4% of lowly methylated genes were core or high copy number genes (Classes A and C), while most highly methylated genes had no known protein domain (Class B, 80.2%). In highly methylated genes, protein domains with the highest representation belonged to transposons (11.9%), high copy number and crinkler gene classes (Class C, 4.8% and crinkler, 0.4%). Core Class A genes therefore tend not to be found in highly methylated regions, consistent with their presence in genomic compartments that are permissive to transcription.

**Figure 3.**
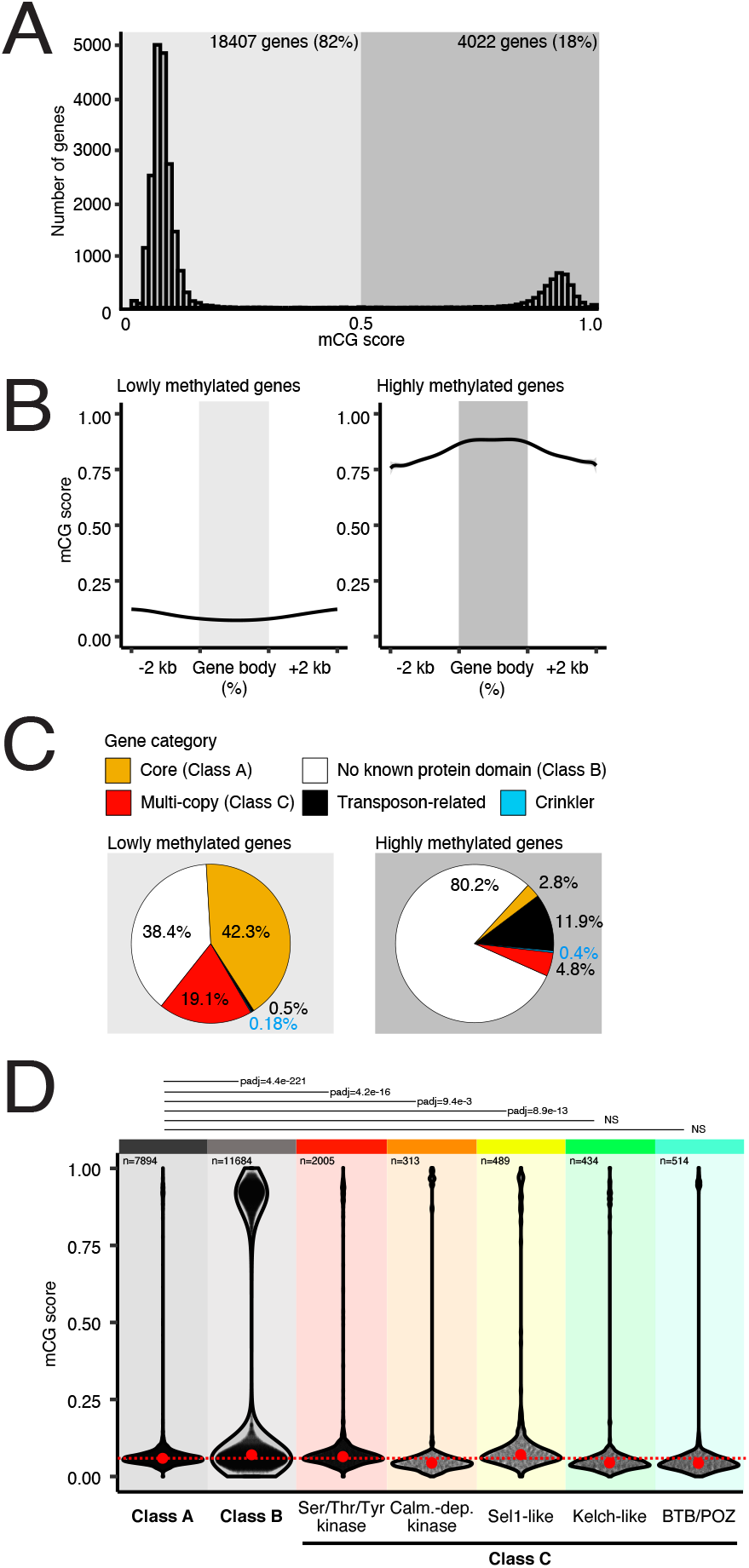
Methylation state and location of genes relative to transposons. Methylation levels of *R. irregularis* genes. **B**. Metagene plots of mCG methylation across genes with high (left) or low (right) types and their 2-kb upstream and downstream sequences. **C**. Protein domain predictions of lowly and highly methylated genes. Pie charts show percentage of genes of each class, as categorized by the identity of their protein domains. Core (class A) are non-repeated genes with an identifiable protein domain. Genes classified as containing no known protein domain (Class B) contain no known protein domain. The repeated gene category includes serine/ threonine/tyrosine kinase, calmodulin-dependent kinase, sel1-repeat, kelch-like and btb-poz domain proteins. Transposon-related genes contain domains such as reverse transcriptase. Crinkler-type genes have a crinkler domain. **D**. mCG score distribution of members of repeated gene families, Class A and Class B genes (grey), and repeated Class C genes (rainbow-coloured). A Kruskal-Wallis Dunn’s multiple comparisons test (Benjamini-Hochberg correction) comparing mCG score distributions of gene groups was used to assess significance. Only adjusted p-values for comparisons to class A are shown.

We then compared the methylation scores of all Class A, B and C genes. Class A and C genes had low global methylation scores, where the latter consisted of a number of sub-groups of genes that each showed different methylation score distributions when compared to Class A. Serine/threonine/tyrosine kinase and Sel1 gene families in particular displayed significantly higher mCG scores than Class A, while other sub-groups either had lower mCG scores or no significant difference (Figure 3D). Strikingly, the CG methylation status of Class B genes displayed a bimodal distribution similar to that of transposons (see Figure 2B). Overall, this data indicates that subsets of Class B and Class C genes are found in highly methylated regions which also contain transposons. On the contrary, Class A genes show a strong tendency to be located in lowly-methylated regions. Differences in the mCG context of gene classes resembles the way some pathogenic fungi genetically compartmentalise their effector genes in TE-rich regions, displaying a so-called two-speed genome (32). Looking at the intergenic distances of genes, we could not find evidence that *R. irregularis* carried a two-speed genome (Figure S3A). However, we observed that class A genes tend to harbour shorter intergenic distances, compared to class B and C genes (Figure S3B-D). These data indicates that high copy number and orphan genes are located in regions that are more gene-sparse, which could be repeated sequences or TEs.

Supporting this observation, high copy number genes (Class C) in *R. irregularis* were previously reported to be localised in proximity to TEs (8). As we found Class B and C genes to be highly methylated and located in gene-sparse genomic regions, we hypothesised that expansions of these gene classes could have been caused by TEs, and wondered if it could be linked to a specific TE family. We first examined how often TEs of each family were the closest element to genes (Table S3). MULEs were the most represented element, consistent with their known bias for inserting near genes (33, 34). For the top three TE families found near genes, we then quantified the frequency at which each element could be found next to members of each gene category, and the distance between them (Figure 4A, bottom and top panel respectively). MULE and CMC/EnSpm elements were slightly over-represented next to Class B genes, compared to Class A genes. All three TE classes were found significantly closer to Class B genes than they were to Class A genes. This indicates that the genomic location and distance of Class A and B genes relative to TEs is different. Class B genes were represented in highly methylated genes (Figure 3C), suggesting a link between their closeness to TEs and methylation status. Gypsy and CMC-EnSpm elements were significantly under-represented next to most sub-groups of Class C genes, compared to Class A. They were either further away from, or located at a distance that was not significantly different to this element’s distance from Class A genes. MULEs however, were the most represented TE family found proximal to all sub-groups of Class C genes and, except for BTB-POZ, they were closer to Class C genes than they were to Class A genes. Thus, Class C genes have a location bias and are often found close to MULE elements specifically. The overrepresentation of MULEs in the proximity of repeated and orphan genes suggests their involvement in expanding these specific gene families. This has has been observed in previous studies of plant genomes in which MULEs are particularly active (35, 36). In rice, MULEs carrying gene fragments or entire genes, called Pack-MULEs, have resulted in the duplication of sequences from over 1500 genes (37). In maize, MULEs often contain receptor protein kinase and calmodulin insertions, similar to the phenomenon observed in the *R. irregularis* genome (38, Figure 4A). In addition, we found MULES within close proximity of eight AGO genes, with five of these genes immediately next to MULEs (Figure 4B). AGO proteins associate with sRNAs such as small interfering RNAs and microRNAs, and function in RNA-based silencing mechanisms. We propose that MULEs may have paradoxically caused the expansion of a pathway that is well-known to suppress TE activity in fungi, plants and animals.

**Figure 4.**
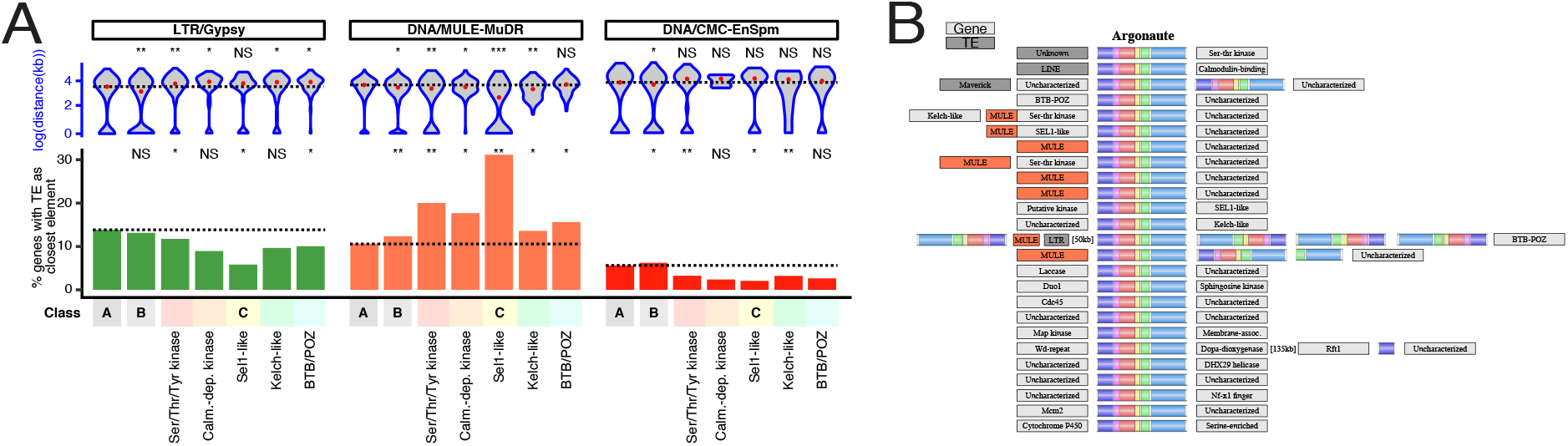
Location of genes relative to TEs. **A**. Identity and distance of TEs closest to genes of classes A, B and C. Underrepresentation or enrichment significance of class B and C genes was assessed by a Fisher’s exact t-test comparing the occurrences of each TE class closest to genes of each family, compared to class A core genes. Top: Log-transformed distance (kb) between genes and closest TE of the displayed classes (Class A, B and C genes, and LTR/Gypsy, DNA/MULE-MuDR, or DNA/CMC-EnSpm TEs). Significance was assessed via a Kruskal-Wallis H test comparing the distance distribution of each TE to repeated gene groups to class A genes. *p-value <0.05 >0.001. **p-value<0.001. **B**. Representation of the genomic context surrounding Argonaute (AGO) genes, a family of repeated genes. The family identity of the closest gene and/or transposable element are shown. MULE elements are indicated in red, alternate TE families are indicated in dark grey, and genes and unknown features are indicated in light grey Coloured regions on Argonautes represent the 6 typical Argonaute protein domains: N-terminal (purple), linker 1 (pink), PAZ (red), linker 2 (yellow), MID (green) and PIWI (blue).

### A subset of *R. irregularis* small RNAs are 2’*-O-*methylated and Argonaute-loaded

As the proximity between MULEs and Argonaute genes suggests a role for TEs in expanding the RNAi gene repertoire, we wondered whether RNAi was involved in the regulation of TEs. RNAi typically relies on four core components: Dicer, Argonaute (AGO), sRNA 2′*-O*- methyltransferase (HEN1) and RNA-dependent RNA polymerase (RdRP) (Figure S4A). sRNAs are generally generated through the cleavage of double-stranded RNA precursors by Dicer proteins and are sorted into specific AGO proteins with different regulatory capacities (39). AGO proteins bind sRNAs and use them as guides to base pair with RNA targets and trigger their repression (40). HEN1 is an RNA methyltransferase that 2′-*O*-methylates the 3′end of sRNAs, protecting them from degradation by exonucleases and increasing their stability (41). In eukaryotes, RdRPs are recruited to target RNAs, which they use as a template to produce complementary RNAs which trigger a secondary amplification mechanism to generate more sRNAs and enhance silencing activity (39). Examination of RNAi pathway genes in the genomes of mycorrhizal fungi revealed that AM species harbour AGO and RdRP gene expansions, and have maintained the sRNA methyltransferase (*HEN1*) gene (Figure S4B, 6, 13, 14). For RNAi pathways to be functional in spores, proteins involved in sRNA biogenesis and function must be expressed. We therefore used label-free quantitative proteomics to profile global protein expression in the *R. irregularis* spore development assay. We first removed any peptide that could be derived from rice root exudates or that matched to multiple proteins, leaving 3475 *R. irregularis* proteins for further analyses (Figure S4C). In all treatments examined, we detected unique peptides mapping to *DCL*, *HEN1*, 10 *AGO* proteins, 3 AGO-binding proteins (*ARB1* and *ARB2* in fission yeast, (42), and 2 genes involved in RNAi-mediated heterochromatin assembly (*HRR1* and *STC1* in fission yeast*, 43, 44*), but found no *RDRP* (Figure S4D, Table S4). Importantly, because our analysis relied on the identification of unique peptide matches, proteins with highly similar sequences (such as *AGO*s and *RDRP*s), could be absent from out dataset but still be expressed. Most RNAi pathway protein components are detected and since *RDRP*s are not always essential for sRNA biogenesis and function, it is reasonable to expect functional RNAi in spores. Detection of *HRR1* and *STC1* homologs suggests the existence of an RNAi-coupled chromatin modification pathway in *R. irregularis*, perhaps in addition to transcriptional or post-transcriptional RNA silencing. Levels of RNAi pathway proteins were relatively high compared to the distribution of label-free quantitation scores for all proteins (Figure S4E). Proteins typically involved in sex and meiosis, as well as putative effectors, were detected (Table S4). We then compared protein expression levels during the spore development assay. Of these, 111 proteins were differentially expressed in at least one condition compared to untreated controls (Figure S4F, Table S5). One AGO protein (A0A2H5UB68) was significantly downregulated at 48h in both mock and exudate treatments, suggesting active regulation of an RNAi factor during spore development. PCA of protein expression indicated that biological replicates do not form discrete clusters based on treatment or time point (Figure S4G). This may indicate subtle protein expression dynamics under these conditions. Functional enrichment of differentially expressed proteins revealed significant GO terms in 24h and 48h exudate treatments that are respectively associated with glycine transport and DNA replication, suggesting active regulation of amino acid metabolism and replication in spores (Figure S4H).

Expression of the sRNA methyltransferase *HEN1* suggests that this enzyme could be actively modifying sRNAs in spores. We set out to investigate the presence of 2′-*O*-methyl modifications of sRNA, and to profile the full spectrum of functional AGO-sRNA complexes present in *R. irregularis* spores. We sequenced and compared the sRNA repertoires sequenced from three RNA extraction methods: 1) total RNA extraction, 2) enrichment of 2′-*O*- methylated sRNAs using sodium periodate (NaIO_4_) oxidation (45), and 3) TraPR column-based isolation of AGO-loaded sRNAs (46)(Figure 5A). We found that NaIO_4_ treatment and TraPR isolation produced similar sRNA profiles with a more well-defined peak than those observed using a total-RNA treatment (Figure 5D–F). The NaIO_4_ and TrAPR sRNA profiles displayed a length distribution centred at 24 nucleotides and a strong bias for sequences beginning with a 5′ terminal uridine or adenine (Figure 5EF). In plants and animals, the identity of the 5’ terminal nucleotide often determines which AGO protein a sRNA is loaded into (47, 48). The 5′U and 5′A biases observed here suggest an evolutionary pressure for sRNAs to begin with a specific nucleotide, perhaps driven by structural specialization of sRNA binding pockets of *R. irregularis* AGO proteins.

**Figure 5.**
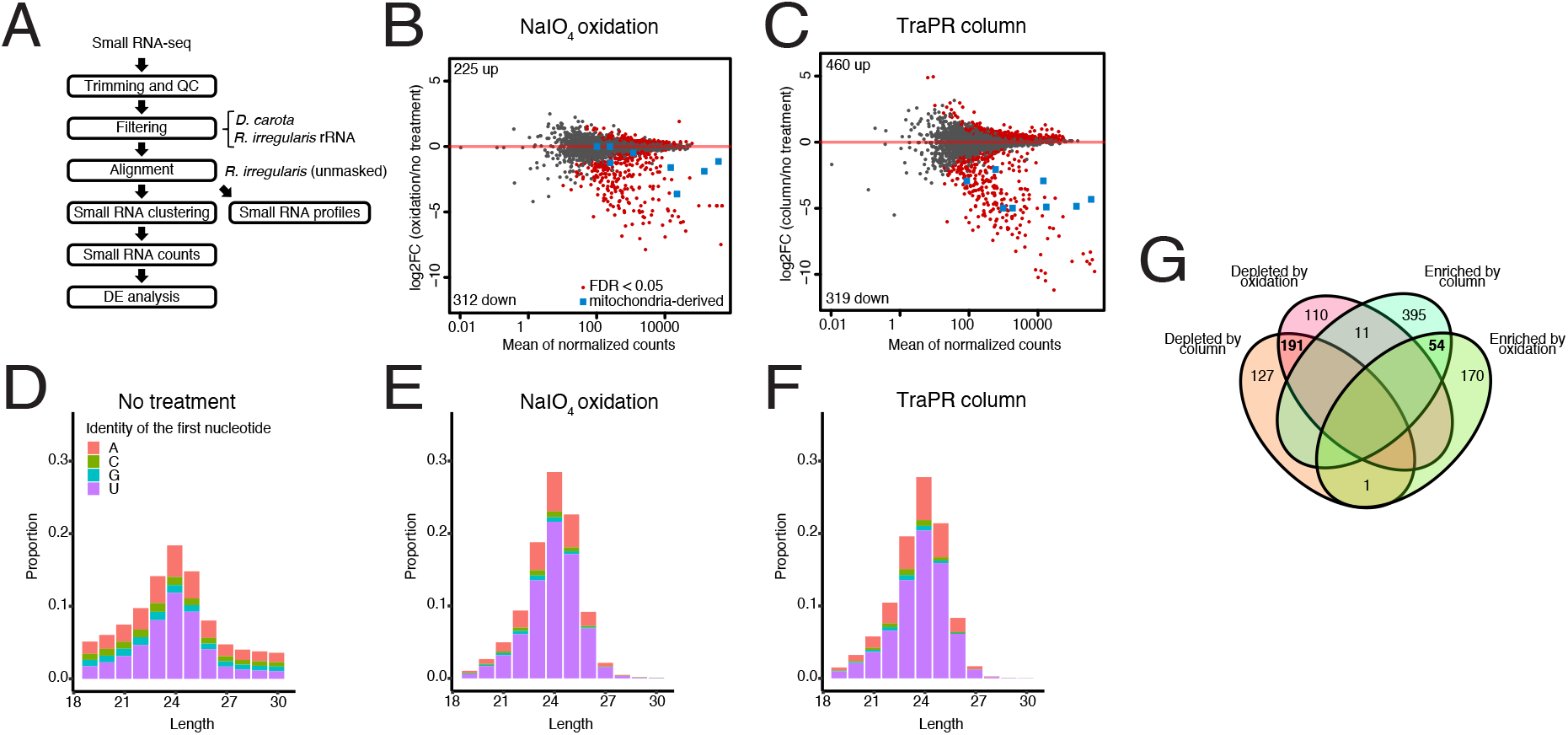
Isolation of 2′*-O-*methylated and Argonaute-loaded small RNA. **A**. Schematic representation of the small RNA-seq analysis pipeline used in this study. Sequence reads were aligned to the *D. carota* and *O. sativa* genomes, and to rRNA sequences matching to *R. irregularis* DAOM 181602=DAOM 197198 on the SILVA database. Reads aligning to any of these were removed and the remaining reads were aligned to the Maeda et al. genome assembly and profiled. Reads were organised into genomic clusters (ShortStack) and counted (FeatureCounts) before differential expression analysis (DESeq2). **B**.**C**. Plot of mean normalized counts against log2 fold change (log2FC) in sRNAs sequenced following enrichment using NaIO_4_ oxidation (B) or TraPR column purification (C), compared to sRNAs sequenced following no treatment. Points represent individual small RNA loci. Significantly differentially expressed loci are shown in red. Mitochondria-derived small RNA loci are shown in blue. **D-F**. Length distribution and first nucleotide bias of the small RNAs in untreated (**D**), NaIO4 treated (**E**) and TraPR column-extracted (**F**) small RNA libraries. **G**. Significantly differentially expressed sRNA loci. sRNAs enriched by NaIO_4_ or column-purification were induced compared to expression in untreated samples and sRNAs depleted by NaIO_4_ or column-purification were downregulated.

We then compared sRNA repertoires produced by the NaIO_4_ and TraPR column treatments. The clustering analysis of sRNA loci revealed that both methods deplete most mitochondria-derived reads. Mitochondria-derived sRNAs were highly abundant, sometimes representing over 10% of all reads. Of 3,495 small RNA loci, 631 were significantly enriched by either column treatment, oxidation treatment or both (Figure 5G). Among those, 54 sRNA loci were enriched by both treatments, pointing to a subset of sRNAs being both 2′-*O*-methylated and Argonaute-loaded. AGO-loaded sRNAs can be expected to be functional and, although the purpose of sRNA modification in AMF is unknown, the 2′-*O*-methylated group likely represents a pool of highly stable sRNAs. sRNA loci significantly depleted or not enriched by either method were removed, leaving a total of 3067 sRNA loci for further analyses. Overall, this data shows that *R. irregularis* spores contain AGO-loaded and 2′-*O*-methylated sRNAs.

### Small RNAs are produced from transposons, methylated regions and their surroundings

To define the genomic origin of sRNA loci, we searched for overlap with genomic features. We found 1510 (49%) sRNA loci matching classified TE sequences, of which 7% showed significant RNA expression (Figure 6A). Of the remaining sRNA loci, 41% were derived from unannotated regions (no gene, no classified TE) and 10% originated from expressed protein-coding genes. No sRNA locus was derived from non-expressed protein-coding genes (data not shown). The majority of genes that produced sRNA had no known protein domain (Class B). sRNA loci derived from non-expressed TE loci were in general highly methylated, while sRNA loci derived from expressed TEs had lower methylation scores (Figure 6B). sRNA loci originating from unannotated regions displayed a bimodal distribution similar to that of TEs and Class B genes (see Figures 2B and 3D). Lastly, coding genes producing sRNAs were mostly non-methylated, consistent with their expression. Copies of all TE superfamilies (DNA, LTR, LINE and Helitron) produced sRNA (Figure 6C). Most sRNA loci were derived from LINE and Gypsy retrotransposons, with respectively 12.6% and 13.5% of elements of each family producing sRNA. sRNA-targeted TEs tended to have a lower divergence (Figure 6C), suggesting a role for sRNA in scanning recently active transposons. The production of 24-nt long sRNA from young TE loci is also observed in plants, where 24-nt mobile sRNAs direct DNA methylation to silence active elements (49, 50, 51).

**Figure 6.**
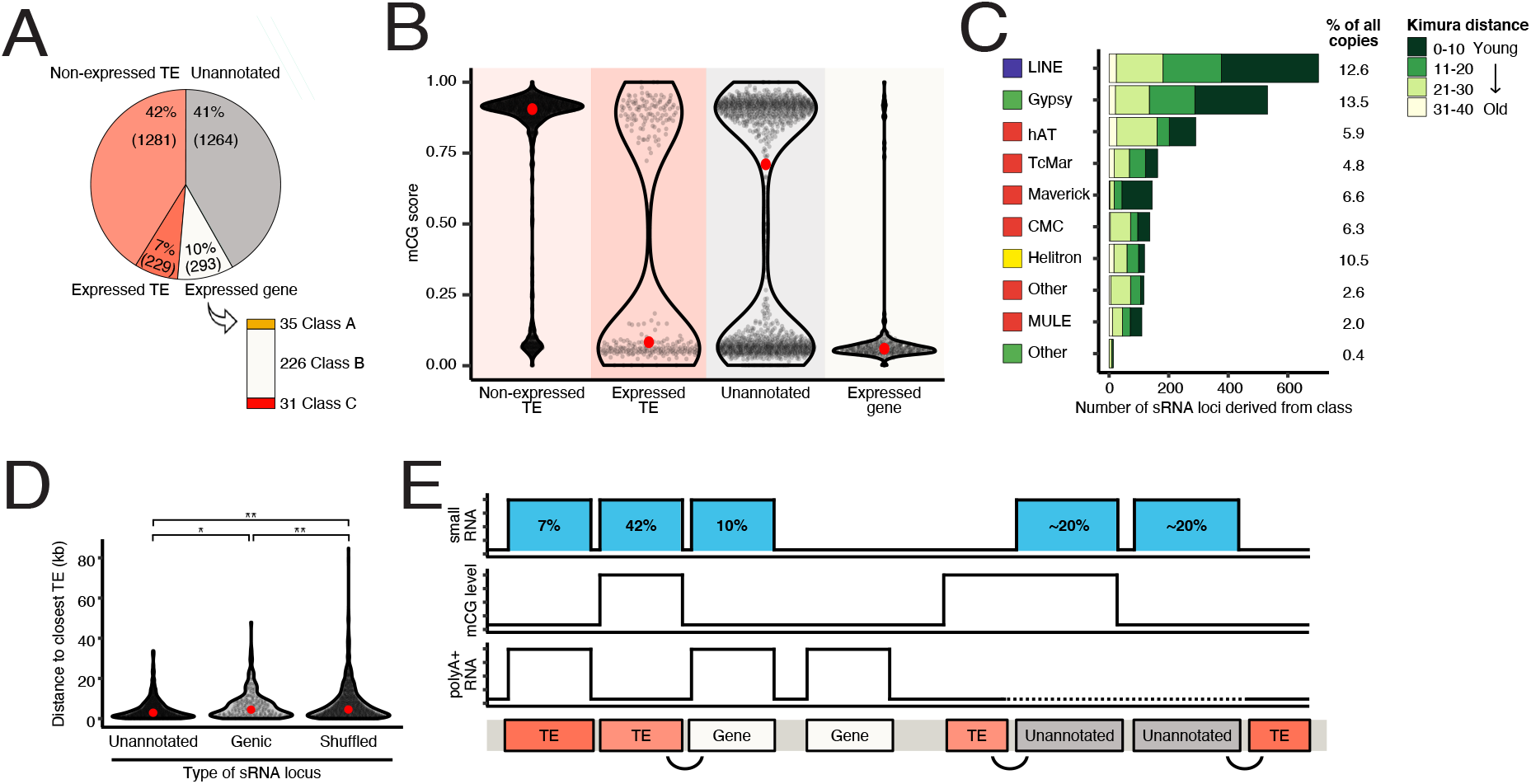
Genomic origin of small RNA-producing loci. **A**. Genomic location of 3067 *R. irregularis* small RNA loci. The number of sRNA loci derived from non-expressed and expressed TEs, unannotated regions and genes is represented in a pie chart. Bar chart shows the number of genes of each class that produce sRNA. **B**. mCG scores of expressed small RNA loci derived from non-expressed TEs, expressed TEs, unannotated features, and expressed genes. Red dots represent the median mCG values of sRNAs from each feature. **C**. Number of sRNA loci associated with different TE superfamilies. Percentage of TE copies from each superfamily that produce sRNA is shown on the right. Green colour code indicates the relative age of sRNA-producing TEs, represented in Kimura distance bins between 0 and 40. Expressed sRNAs have been grouped into bins based on Kimura distance and hence relative age, represented using colour coding. **D**. Distance from small RNA loci derived from unannotated regions and expressed genes to closest known TE. Significance was assessed by a non-parametric Kruskal-Wallis test comparing the distance distribution of unannotated or genic sRNA loci to shuffled loci respectively. A Kruskal-Wallis H test comparing unannotated and genic sRNA loci to shuffled loci was used to assess significance. * p = 2.8E-06, **p<2.2E-16. **E**. Schematic diagram of the genomic distribution of small RNA loci hypothesised due to data from this study of the *R. irregularis* genome, sRNAome, methylome and transcriptome. sRNA loci, methylation levels and RNA expression are displayed. The proportion of sRNA loci corresponding to each genomic context is indicated in blue boxes. Curvy lines highlight the proximity between TEs and genic and unannotated sRNA loci. Expression of unannotated regions was not analysed and is represented by a dotted line.

We next asked what could explain the expression of sRNA from non-TE regions. We observed that sRNA loci derived from unannotated regions and genes were located significantly closer to TEs than a set of simulated sRNA loci shuffled randomly throughout the genome. While we do not know what the function of these loci are or why these specific genes produce sRNA, their proximity to TEs suggests a link with TE activity or regulation. Together, our data shows that most small RNAs are produced from TEs (49%), unannotated regions (∼20%) or genes in the vicinity of TEs (∼20%) (Figure 6E), indicating a potential role of sRNA in regulating TEs.

## Discussion

### Signs of recent or ongoing TE activity in *R. irregularis*

Transposons are major drivers of genome evolution due to their activity as powerful mutagens and have the potential to inflate genome size and physically move DNA to new locations within the genome. Pathogenic fungi associated with a range of plant hosts have an abundance of TEs, thought to facilitate adaptability and enable these species to cope with varied ecological niches and diverse host species (52, 53, 54). Compared to other fungal lineages, fungi that are tightly linked to plants have been noted to possess larger genomes, and this can be explained by their repeat content (53, 54). Genomes of AM fungi also have high repeat contents (this study, (8, 55), but we could only classify 12% of the genome sequence into TE families. 8% of repeats consist of high copy number and orphan gene families, and 24% of repeats are of unknown origin. We found evidence of expression for TEs belonging to all superfamilies. Moreover, 39 TE subfamilies (e.g. MULE-MuDR and Gypsy) are dynamically regulated during spore development, a potential indicator of active mobilization and ongoing epigenetic regulation.

We provide two lines of evidence of epigenetic regulation occurring in *R. irregularis: 1*)high CG methylation levels at the majority of TE loci, and 2) sRNA production at some TE loci, particularly evolutionarily young TEs. Consistent with the activity of these pathways in spores, we detect protein expression of a DNMT1-related DNA methyltransferase (A0A2H5UGI7, Table S4) and 17 proteins typically involved in sRNA biogenesis and function (Figure S4D). Interestingly, we found examples of expressed TEs that were lowly methylated, suggesting a correlation between relaxation of epigenetic silencing and TE expression. There is therefore potential for epigenetic mechanisms and transposon activity to be actively shaping the genome of *R. irregularis*. In AM fungi, defence against TEs likely involves DNA methylation and RNAi. The mechanism(s) by which sRNAs act in *R. irregularis* remain to be investigated, and could include RNA-dependent DNA methylation, histone modification, transcriptional silencing and/or post-transcriptional silencing.

### The non-core gene repertoire of *R. irregularis* may be linked to TE activity

Annotation of AM fungal genes revealed a larger gene repertoire than other species of fungi (56, 57). Intriguingly, 45% of *R. irregularis* genes have no known protein domains, and are of unknown origin and function (Class B genes in this study, and reviewed in 5). Similar to other AM fungi, 16% of *R. irregularis* genes are members of expanded families such as protein kinases (Class C). We now find that Class B and Class C serine/threonine/ tyrosine kinases and Sel1-like genes tend to be more highly methylated than core genes (Class A) and to be located in gene-sparse genomic regions. Although this genomic compartmentalisation is not as well defined as that observed in the two-speed genomes of pathogenic fungi, our data suggests that core and repetitive genes are not evenly distributed throughout the genome. A sister phylogenetic lineage of AM fungi, *Geosiphon pyriformis* (58), displays a non-compartmentalised genome similar to what we observe in *R. irregularis*, although the comparison of intergenic distances between different classes of genes remains to be assessed.

In fungi, limited methylation occurs in gene bodies (18). In *R. irregularis*, we found that 18% of genes are highly methylated and these genes are mostly orphan sequences found near TEs. Because of their repetitiveness, high methylation levels, proximity to TEs and unknown evolutionary origin, some of these genes could alternatively be previously uncharacterised TEs. Future work in TE discovery and classification, and subsequent improvements in the annotation of AM genomes may shed light on the origin of these sequences.

### Proximity between MULEs and high copy number genes suggests transposon-mediated expansions

Genomes analyses of five *R. irregularis* isolates revealed that roughly half of their gene repertoires is shared (5). Most unshared genes belong to Class B and C (orphan and high copy number), raising questions about their evolutionary origins and biological functions. As we find these genes to be located in gene-sparse regions that are likely repetitive, copy number variations could be explained by difficulty assembling genomic repeat sequences. Chromosome-level genome assemblies of *R. irregularis* isolates and AMF species should clarify the origin of these variations. Nevertheless, the proximity between TEs and specific gene families suggests a role for TEs in causing gene expansions. We found that MULEs were more often found next to Class B and C genes, in comparison to Class A genes which did not show this tendency (or which were more randomly distributed in their proximity to MULEs). We therefore propose that gene expansions in *R. irregularis* may have been caused by MULEs. As two subfamilies of MULEs are differentially expressed during spore development, recent or ongoing MULE activity could explain some of the genomic variations observed in different *R. irregularis* isolates. Interestingly, expansion of Argonaute genes also seems to be linked to MULEs, suggesting that RNAi pathway(s) may have been shaped by TE activity in AM fungi.

### Adaptation and evolution driven by TE activity

Perhaps the most intriguing feature of AM fungi is their thus far undocumented sexual reproduction. While there appears to be potential for them to reproduce sexually (59, 60, 61, 62), meiotic events have never been observed and genetically distinct strains created by meiotic divisions have never been identified. Nevertheless, sexual or parasexual reproduction could exist in AM fungi. In the absence or rare occurrence of sex, or perhaps in parallel to sex, three mechanisms could generate genetic variation: 1) horizontal gene transfer, which is known to occur in AM genomes (63, 64), 2) TE activity, and 3) cryptic recombination, both of which were proposed to occur in AM fungi (5, 65). One could predict that these mechanisms bear more important roles in genome evolution of asexual than sexual species. TE activity generates significant adaptive genetic variation and has been shown to play roles in the evolution of genes encoding proteins involved in host interaction (66). Because TE-linked orphan and high copy number genes constitute the majority of *R. irregularis* genes, an important question arises: do these genes have roles in plant-AMF interactions, or are they just by-products of TE movement? Are multi-copy genes with no known protein domain functional genes, or are they unclassified TEs/repeats? We propose that among all expanded genes, AGOs are the genes most likely to promote fitness, by controlling TE activity. As *R. irregularis* sRNAs appear to target TEs and regions located near them, we speculate that AGO expansions may help to strike a balance in the evolutionary conflict between TE invasion and TE-driven adaptation.

As TE mobilisation through a population is thought to be facilitated by sexual reproduction of their hosts, can TEs be strong drivers of genome evolution in asexual organisms? Interestingly, bdelloid rotifers are asexual animals that display features similar to those observed in *R. irregularis*: 1) recent and ongoing TE activity, 2) large expansions of RNAi pathway genes (∼ 22 Argonaute, 4 Dicer and 37 RdRP) (67), and (3) targeting of TEs by small RNA (68). In this case, cryptic recombination was not found to support the inheritance of TE loci. While the cause of rotifer RNAi gene expansions is unknown, this research provides a case study of concomitant TE activity and expansion of RNAi genes in an asexual species. Without sex and recombination, organisms have limited ways of defending their genomes against unchecked TE expansions. As Nowell et al., propose (67), RNAi may have been required for ancient transitions from sexuality to asexuality. Thus, a controlled balance between TE activity and silencing may have enabled the long term ecological success of AM fungi. Future analyses of high-quality genome sequences and annotations are required to shed light the genomic organisation and evolution of AM fungi.

## Methods

### Production of rice exudates

The rice (*Oryza sativa subsp. janponica*) cultivar Nipponbare was used to produce exudates for this study. Rice seeds were manually de-husked and sterilised by incubating with 3% sodium hypochlorite for 30 minutes on a platform shaker, followed by rinsing three times with diH_2_O. Sterile seeds were subsequently pre-germinated on 0.7% Bacto Agar plates for four days at 30°C. Seedlings were then transferred to trays of autoclaved sand and grown for six weeks in a growth chamber under a 12-hour light/12-hour dark cycle at a 28°C daytime/23°C nighttime temperature and 60% humidity. From two weeks onward, rice plants were provided twice a week with half-strength Hoagland’s solution (25 μM phosphate) containing 0.01% (w/v) Sequestrine Rapid (Syngenta). For exudate collection, the six week-old plants were removed from soil, rinsed, and placed into 250mL conical flasks containing 200mL half-strength Hoagland’s solution, with the roots submerged. Three rice plants were placed into each flask, which were then incubated for three days at the previously-described plant growth conditions on a mechanical shaker at 50 shakes/min. Following the 3-day incubation, the half-strength Hoagland’s solution containing plant exudates was filter-sterilised using 0.2μM filters. Half-strength Hoagland’s solution was prepared for use as a control treatment, and was incubated without the addition of rice plants for three days at plant growth conditions before subsequent filter-sterilisation.

### R. irregularis in vitro spore development assay

*R. irregularis* DAOM 197198 grade A spores (Agronutrition, Toulouse, France) were suspended in 1x M Media at a concentration of 10,000 spores/mL. For a total of 20 samples, 5mL (50,000 spores/sample) of the spore solution was aliquoted to individual 16.8mL tissue culture wells, and the samples were incubated for seven days at 30°C and 2% CO_2._ Following incubation, spore samples were either immediately frozen in liquid nitrogen for a 0h time point or were re-incubated at 30°C and 2% CO_2_ with either half-strength Hoagland’s solution or with sterilised rice exudates, for 24h or 48h hours. Four spore samples were produced per treatment. Prior to freezing, all spore samples were drained using 40μM cell strainers. For use in downstream processing, liquid-nitrogen frozen *R. irregularis* spore samples were homogenised using a mixer mill MM 400 and 25ml grinding jars (Retsch), shaking at 25 shakes/second for 20 seconds.

### HMW DNA extraction and sequencing

High-molecular weight DNA was extracted using the protocol from Schwessinger & McDonald (69). 100mg of ground spore material was resuspended in lysis buffer and processed as indicated. Two successive rounds of cleanup were performed using a 0.45X volume of Ampure XP beads in DNA-Lo-Bind tubes following the Manu^-^facturer’s protocol. DNA was finally eluted in 50uL of 10mM Tris-pH8. DNA quality was assessed by running on a 0.5% agarose gel. Sequencing libraries were prepared using the Oxford Nanopore Rapid DNA sequencing kit SQK-RAD004 and sequenced on MinION flow cells FLO-MIN106D following the accompanying protocol.

### RNA extraction, library preparations, and small RNA treatments

RNA extraction using an RNeasy Plant Kit (Quiagen, Germany) was carried out on a portion of each ground spore sample. RNA integrity and purity were assessed using an Agilent 2100 Bioanalyser and RNA 6000 Pico Kit (Agilent, USA) and a Tapestation (Agilent). Paired-end polyA+ RNA-libraries were produced and sequenced by Novogene UK Co. Ltd. with read lengths of 150 bp. Small RNA libraries were produced using the NEXTFLEX® Small RNA-Seq Kit v3 following the gel-based protocol. Small RNA libraries were sequenced on an Illumina Illumina HiSeq 1500 with read lengths of 50 bp.

For small RNA profiling experiments, 15-20mg of the ground spore samples were then split into three equal volumes and subjected to one of three treatments: NaIO_4_ oxidation, TRaPR column treatment, and a non-treated condition. For the oxidation treatment, 5uL of 200mM NaIO4 were added to 500ng total RNA diluted in 24.5 uL of 1X borate buffer. Oxidation was performed at room temperature for 10 minutes, then RNA was precipitated at −20 for 1h with 0.1V (4 uL) 3M Sodium acetate and 2.5V(150uL) ice cold 100% ethanol. RNA was centifuged at 13000RPM at 4degC for 20 mins, pellets were washed twice with 0.2ml ice cold 80% ethanol, spun at 4degC for 5 mins, air dried for 10 mins, resuspended in 10.5 nuclease-free H20, then ligated following the NEXTFLEX Small RNA-Seq kit protocol. For Lexogen’s TraPR™ Small RNA Isolation Kit, 20mg of ground spore material was suspended in TRaPR lysis buffer and the standard TRaPR experimental procedure and RNA extraction was carried out (Lexogen, Austria). The non-treated ground material of each sample was also suspended in TRaPR lysis buffer and RNA extracted through a phenol-chloroform extraction. All treated and RNA-extracted samples were then used to produce small libraries for sequencing using a NETFLEX small RNA-Seq kit (PerkinElmer, USA).

### Proteomics

20 mg of each ground spore sample was resuspended in 1x LDS Buffer and 100 mM DTT and incubated at 70°C for 10 min. Proteins were separated on a 10 % NuPage NOVEX Bis-Tris gel (Thermo) for 8 min at 180 V in 1 × MES buffer (Thermo). The gels were fixated, stained with Coomassie Brilliant Blue G250 (Sigma) and after^-^wards destained with water. In-gel digestion and desalting on C18 StageTips were performed as previously described (70, 71). LC-MS/MS analysis was carried out on an EASY-nLC 1000 system (Thermo) coupled to a Q Exactive Plus Orbitrap mass spectrometer (Thermo) via the nanoflex electrospray ion source. Peptides were separated on a 25 cm reversed-phase capillary with a 75 μm inner diameter packed in-house with Reprosil C18 resin (Dr. Maisch GmbH). The peptides were eluted during a 208 min gradient from 2 to 40 % acetronitrile in 0.1% formic acid at a constant flow rate of 225 nl/min. The Q Exactive Plus was operated with a top 10 data-dependent acquisition method. For raw file peak extraction and the identification of protein groups the MS raw files were searched with MaxQuant (version 1.6.10.43; (71) against the following three databases from UniProt: UP-000236242 (*Rhizophagus irregularis*), UP000059680 (*O. sativa subsp. japonica*) and UP000007305 (*Zea mays*). The database searches were performed with MaxQuant standard settings with additional protein quantification using the label free quantification (LFQ) algorithm (72, 73) and the match between runs option was activated. The data was further analyzed in R (version 3.6.2) using an in-house script. In short, from the identified protein groups known contaminants, reverse entries, protein groups only identified by site or with no unique or less than two peptides were filtered out and excluded from the analysis. Missing LFQ values were imputed at the lower end of values within each sample and data plotted using the ggplot2 and pheatmap packages (74, 75).

### Annotation of transposable elements

For annotation of transposable elements, the long read PacBio assembly http://nekko.nibb.ac.jp of the *R. irregularis* genome was used. Sequencing with long reads can improve detection of transposable elements, as short read data tend to collapse highly repetitive regions of the genome. *De novo* annotation was performed using RepeatModeler2 76), which uses the RepeatScout (77) and RECON (78) algorithms for TE discovery. We also used LTR_retriever (79) and LTR_harvest (80) to enhance detection of LTR retrotransposons. This combined new library of denovo elements was used to annotate the *R. irregularis* DAOM 197198 genome (8) with Repeat- Masker (81). The age of each TE copy was calculated as % of divergence (Kimura distance) from the consensus models, which were generated by RepeatModeler (76). Transposon families were ‘binned’ according to divergence and plotted in relation to genome coverage (RepeatLandscape). To achieve retrieval of complete copies the consensus sequences for the CMC-EnSpm, Crypton-A and MULE families were manually curated. For this we performed multiple rounds of curation using BLAST (82) to retrieve the top 50 hits for each consensus on the genome, which were then extended, aligned with MUSCLE (83) and manually edited until the complete sequence of the consensus sequence was retrieved.

### Generation of transcriptome data for gene annotation

Material used to produce RNA for gene annotation was treated as follows. *R. irregularis* DAOM 197198 spores were harvested from carrot root organ cultures under sterile conditions by dissolving the phytagel in 10 mM citrate buffer at pH 6.0. Carrot roots were carefully removed to avoid any contamination. Two-week-old *N. benthamiana* seedlings grown in vitro were transferred to autoclaved silver sand and inoculated with 3200 freshly extracted *R. irregularis* spores or water for mock controls. After three weeks, plants were pulled out of sand and roots from four plants were pooled together per biological replicate for both mock and mycorrhized conditions. For germinated spore samples, 60 000 spores were germinated and grown in liquid M-medium in the dark at 30°C supplemented with 2% CO2 for 7 days. Germinated spores were harvested using a 40 μm cell strainer (Sigma) and the excess of M-medium drained out before snap-freezing the samples in liquid nitrogen. All samples were produced in triplicates. Total RNA was extracted as previously described (84). Complementary DNA (cDNA) libraries were prepared from 1 μg RNA using TrueSeq RNA Sample Prep Kit (Illumina, California, USA), according to the manufacturer’s instructions. RNA and DNA libraries were quantified using a Qubit fluorometer (Thermo Fisher Scientific, Massachusetts, USA), and their integrity was checked on a TapeStation 2200 (Agilent Technologies, California, USA) using RNA ScreenTape (Agilent 5067-5576) and High Sensitivity D1000 Screen Tapes (Agilent 5067-5584), respectively. Libraries were diluted to 4 nM and sequenced on a NextSeq 500 Sequencing System (Illumina, California, USA) using NextSeq 500/550 High Output Kit v2 (150 cycles, paired-end 75 bp), according to Illumina’s instructions. A maximum of 400 M reads was allocated for colonised *Nicotiana benthamiana* root samples, while samples corresponding to *Rhizophagus irregularis* spores and non-colonised (mock) roots were run together and shared 400 M reads.

### Annotation of genes

Gene models were predicted using the BRAKER2 pipeline (85) on the contigs generated by Maeda *et al*. (8) using transcriptomic data as extrinsic evidence, yielding 22,338 gene models. Gene models’ completeness was assessed by predicting BUSCO genes (86). Functional annotations were lifted from Maeda et al’s gene annota^-^tion by intersecting genes with >50% matching sequences. Genes with transposon-related protein domains were removed (transposon|zinc_finger_bed_domain|ricesleeper|helicase-primase|helicase/primase|gag-pol|far1-related|ribonuclease_hi|ribonuclease_h|jockey|rve_super_family_integrase|transposase|transposable|helitron|pif1| zinc_finger_mym-type_protein_2|reverse_transcriptase).

### Small RNA sequencing data processing

Raw sequencing reads were quality-trimmed using cutadapt 1.9.1 (87) using the parameters recommended in the NEXTflex Small RNA instructions to trim 3’ adapter (-a TGGAATTCTCGGGTGCCAAGG --minimum-length 10), and to trim 4 bases from either side of each read (-u 4 -u -4). Read quality was assessed using FastQC v0.11.4 (88). Clean reads were aligned to a ribosomal RNA library made using SILVA database sequences for R. irregularis, and the Daucus_carota_388.v2.0 host genome using Bowtie-1.2.2 (89) with the following parameters -q -a -v 0. All reads perfectly matching either rRNA or the carrot genome were filtered out. Remaining reads were aligned to the unmasked R. irregularis DAOM-181602 genome using Bowtie-1.2.2 and the following parameters: -q -k 500 -m 50 -v 1. Small RNA profiles were plotted for collapsed reads using custom scripts and the ggplot package. Reads from untreated, oxidised and column-purified libraries were concatenated and clustered using Shortstack using the following parameters: --dicermin 20 --dicermax 27 --foldsize 300 --pad 200 --mincov 10.0rpmm --strand_cutoff 0.8 (90). Small RNA counts were generated using featureCounts 1.5.0 using the Shortstack output following parameters: -M --fraction -T 8 -F GTF -g ID -t nc_RNA (91). Small RNA counts were analysed and plotted using the DESeq2, ggplot2 and pheatmap packages (74, 75).

### PolyA+ RNA sequencing data processing

RNA-sequencing reads were filtered by Novogene to remove low-quality reads (reads containing Qscore <=5 in over 50% of bases, reads containing N > 10%). Reads were also trimmed by Novogene prior to our analysis (5’ adapter - 5′-AATGATACGGCGACCACCGAGATCTACACTCTTTCCCTACACGACGCTCTTCCGATCT-3′, 3′ a d a p t e r - 5 ‘ - GATCGGAAGAGCACACGTCTGAACTCCAGTCACATCACGATCTCGTATGCCGTCTTCTGCTTG-3′). We assessed read quality was assessed using FastQC v0.11.4 (85), all samples showed a Phred score higher than 30. Clean reads were aligned to the unmasked R. irregularis DAOM-181062 genome using STAR 2.5.4 (92). For TE subfamily expression analyses, reads were aligned using the options --outFilterMultimapNmax 100 and --winAnchorMultimapNmax 100 and the counts were generated using the TEtranscripts package with the gene annotation and a curated version of TE annotation files produced in this study (options --mode multi and --stranded no) (93). Our TE annotation was curated to exclude simple repeats, unclassified repeats, low complexity repeats, satellites, rRNAs, snRNAs and tRNAs. TE subfamilies with at least 100 normalised counts were considered expressed. The output count file was used to perform differential expression analysis of TEs with at least 1 normalised count using DESeq2, with 0h untreated samples as a control. TE subfamilies with a P adjusted value below 0.05 and an absolute log2 fold change higher than 0.5 were considered significantly differentially expressed. For locus-level TE expression, counts were generated with featureCounts (subread package 2.0.1, -- fraction parameter used) and only TEs of a length > 100bp and at least 1RPKM in at least 6 samples were considered. For gene expression analysis, reads were mapped using STAR with the option --outFilterMultimapNmax 20 and counts were generated with featureCounts without the fraction option. Genes with at least 2 normalised counts in at least 2 samples were considered expressed. TE and gene counts were analysed and plotted using the DESeq2 ggplot2 and pheatmap packages in R (74, 75).

### ONT reads processing and DNA methylation analysis

Genomic CpG methylation data was produced from Nanopore sequencing data with DeepSignal (0.1.8), called against model.CpG.R9.4_1D.human_hx1.bn17.sn360.v0.1.7+ using default parameters (94). From the DeepSignal output, data for symmetrical CG sites was merged, and overlapped with TE or gene loci using bedtools map - median. Plots were generated using ggplot2 74).

## Data availability

Sequences obtained in this study have been deposited at the NCBI GEO database under accession GSE172187 and at SRA PRJNA722386. Proteomics data are available via ProteomeXchange with identifier PXD025245.

## Acknowledgements

We would like to thank all members of the Miska, Paszkowski, Butter and Schornack laboratories, and Martin Simard for providing valuable input on the manuscript. This work was supported in whole or in part by Cancer Research UK (C13474/A18583, C6946/A14492) and the Wellcome Trust (219475/Z/19/Z, 092096/Z/10/Z) to E.A.M. Research in U.P lab was supported by the Engineering the Nitrogen Symbiosis for Africa (ENSA) project, which is funded by a grant to the University of Cambridge by the Bill & Melinda Gates Foundation. Research in the S.S. lab was supported by the Gatsby Charitable Foundation (GAT3395/GLD) and the Royal Society (UF110073 and UF160413). A.D. was supported by a Natural Sciences and Engineering Research Council of Canada postdoctoral fellowship (PDF-532905-19). IB, was supported by Wellcome grant WT207492 and 104640/ Z/14/Z, 092096/Z/10/Z. For the purpose of Open Access, the author has applied a CC BY public copyright licence to any Author Accepted Manuscript version arising from this submission.

## Supplementary material

Table S1. Differentially expressed TE subfamilies

Table S2. Expressed genes containing TEs or TE fragments

Table S3. Top 100 TEs closest to genes

Table S4. *R. irregularis* proteins detected by mass spectrometry

Table S5. Differentially expressed *R. irregularis* proteins

**Figure S1.**
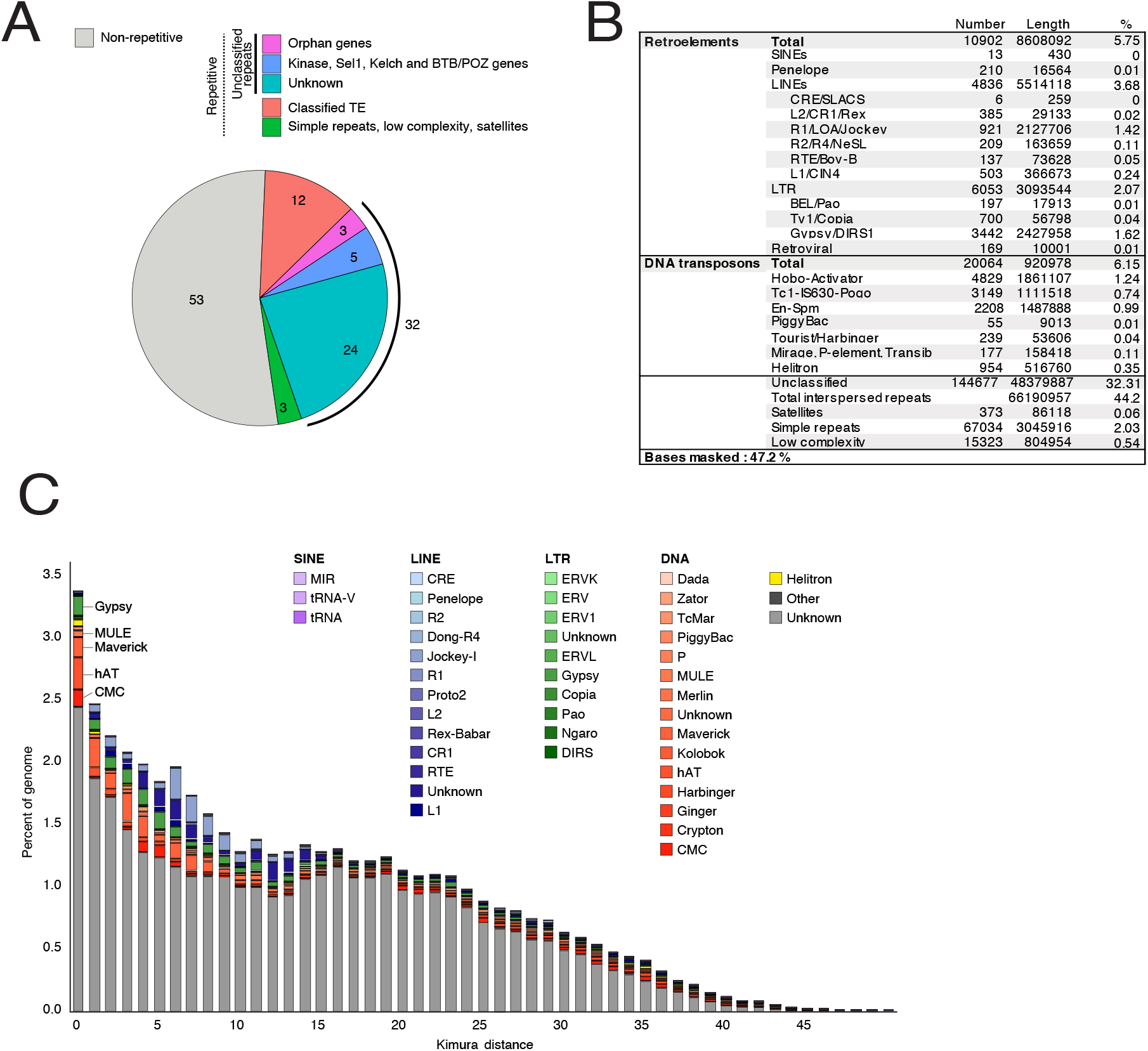
Transposon and repeat annotation. **A**. Summary of genome sequence composition (%). **B**. The composition (%) of major TE families in *R. irregularis*, alongside associated length and copy number. **C**. Kimura distance-based copy divergence analyses of transposable elements in *R. irregularis*. Graph represents genome coverage (%) for each TE superfamily plotted against Kimura distances of TEs (CpG adjusted *K-value* from 0 to 50). TE copies with a low Kimura distance value have a low divergence from the consensus sequence and may correspond to recent replication events. Sequences with a higher Kimura distance value corresponded to older divergence. Data displayed is the same as data used to produce Figure 1A, and also includes unknown/unclassified elements.

**Figure S2.**
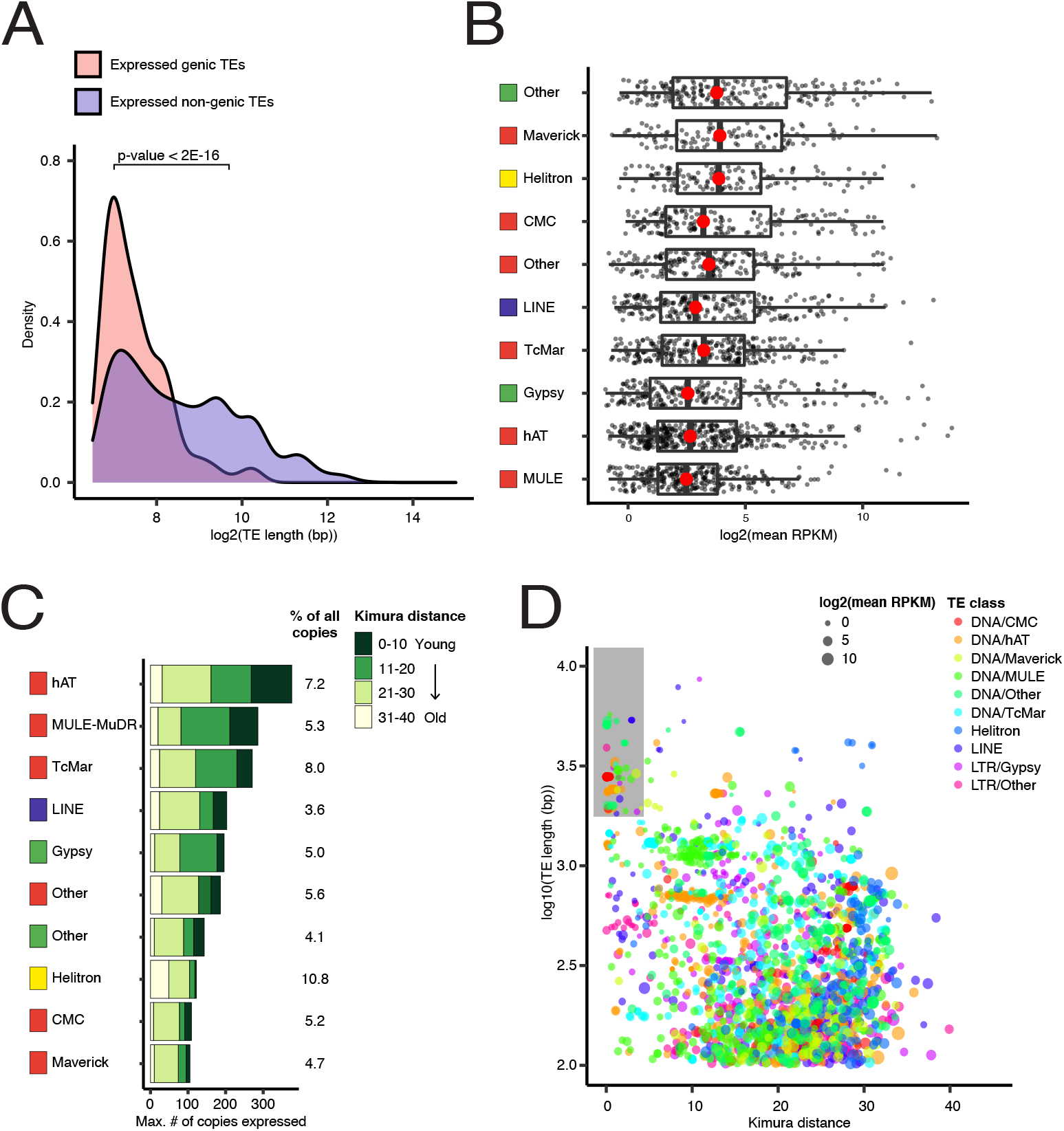
Transposon expression at individual loci. **A**. Density plot depicting log-transformed TE length of genic (pink) and non-genic (purple) expressed individual TE loci. Expressed TEs were classified as ‘genic’ when found overlapping with the coding region of expressed genes. Expressed TEs that did not overlap with the coding regions of expressed genes were classified as non-genic. **B**. Non-genic TE expression levels, grouped by superfamily. Boxplots represent interquartile ranges and red dots represent the medians of log2 transformed mean RPKM across 20 samples (5 conditions, 4 replicates/condition). **C**. Number of expressed non-genic transposon copies of each super-family. Absolute TE numbers are displayed as percentage of expressed TEs compared to all TEs in the genome. Expressed TEs have been grouped into bins based on Kimura distance and hence relative age, represented using colour coding. **D**. Log-transformed length of all expressed TEs relative to divergence expressed as Kimura distance. Colour and point size indicate the TE class and log-transformed mean RPKM values respectively. Shaded area highlights TEs of >2kb length and <5% Kimura distance.

**Figure S3.**
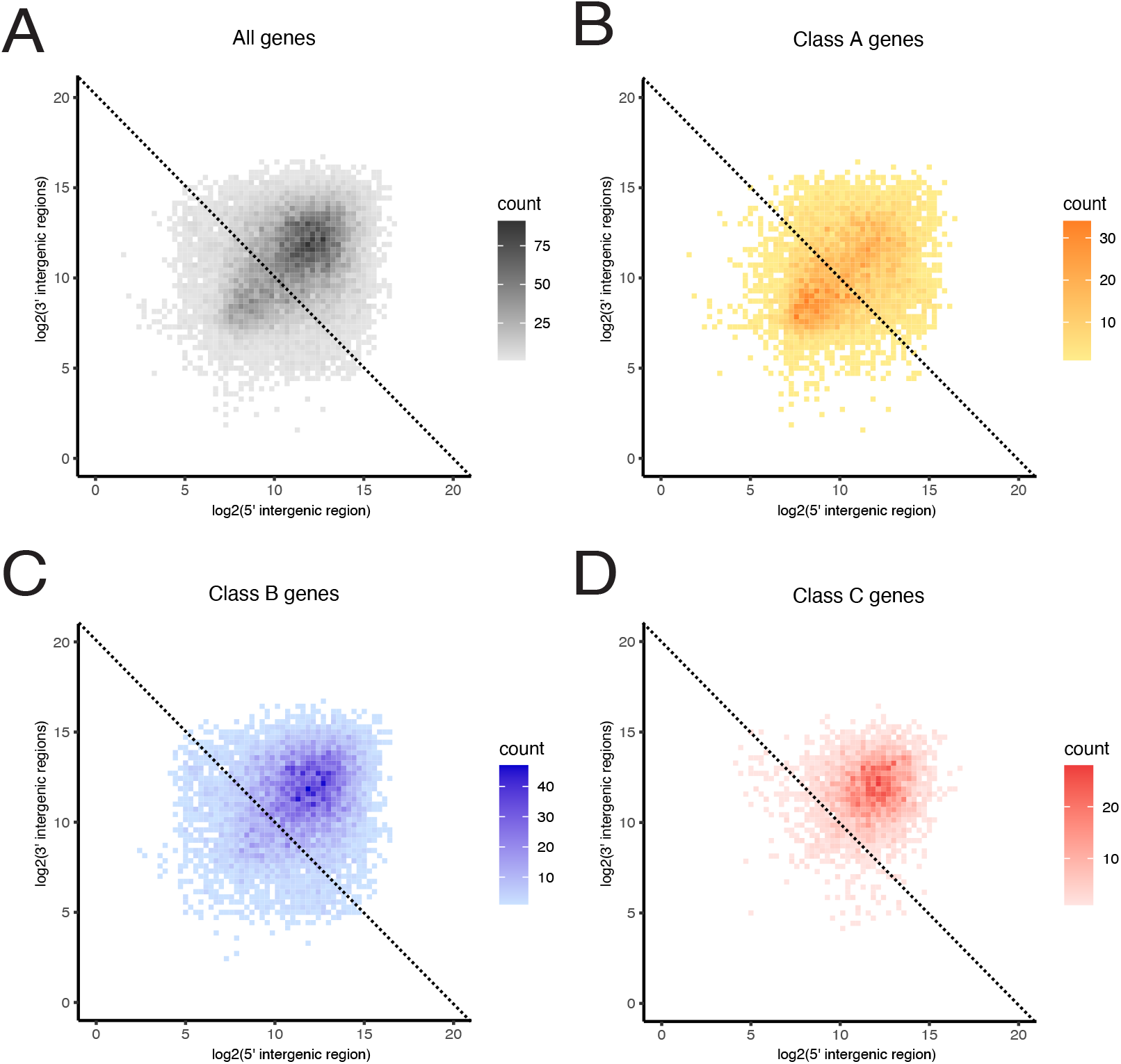
Uneven distribution of genes in the genome of *R. irregularis*. Log-transformed intergenic disances upstream genes (5′) is plotted against log-tranformed intergenic distances downstream genes (3′) for all genes (**A**), class A (**B**), class B (**C**) and class C (**D**) genes.

**Figure S4.**
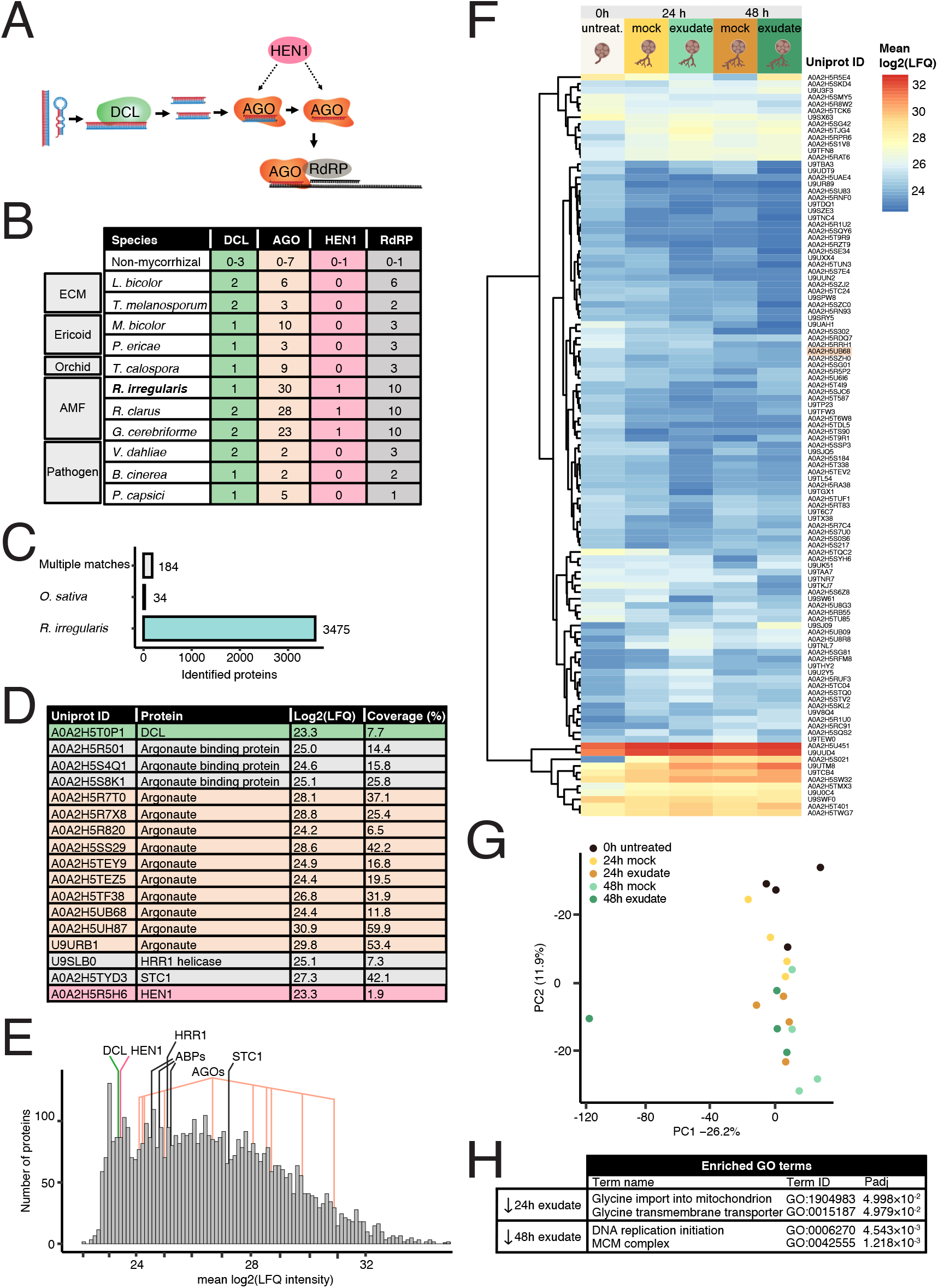
RNAi-related genes in *R. irregularis* and their expression. **A**. Schematic representation of a typical RNAi pathway. Double-stranded RNA or hairpin RNAs are cleaved by the RNAse Dicer, generating sRNA duplexes. HEN1 methylates either the duplex or single-stranded sRNA loaded into AGO proteins. The sRNA-AGO complex then targets RNAs by base pairing. In some cases, RNA-dependent RNA polymerases (RdRPs) are recruited to facilitate silencing by using the target RNA as a template to generate more sRNA. **B**. Number of putative DCL, AGO, HEN1 and RdRP genes found in genomes of species of mycorrhizal species: ectomycorrhizal (ECM), ericoid mycorrhiza, orchid mycorrhiza, arbuscular mycorrhiza (AMF) and pathogenic fungi capable of cross-kingdom sRNA transfer. **C**. Number of proteins identified by mass spectrometry matching to the proteomes of *R. irregularis*, *O. sativa* or both. **D**. Uniprot IDs, label-free quantitation (mean log-transformed LFQ intensities) and % unique coverage of RNAi pathway genes detected by proteomics. **E**. Distribution of LFQ intensity of proteins detected by mass spectrometry. Values for RNAi pathway proteins are indicated. **F**. Heatmap and hierarchical clustering of mean log-transformed LFQ intensities of differentially expressed *R. irregularis* proteins (|Δ log2(LFQ)| > 1.1; log10(Padj) > 1.1) at 24h or 48h post-treatment with mock (Hoagland’s; nutrient condition) or exudate (rice root exudates) treatment relative to control conditions (0h; no treatment). ID marked in orange is an Argonaute protein. **G**. Principal component analysis of protein expression across all replicates and treatments, 0h control, 24h mock and rice exudate treatments, and 48h mock and rice exudate treatment (20 samples total, at 4 replicates per treatment). **H**. Functional enrichment analysis of differentially expressed proteins using g:Profiler.

